# An open-source software analysis package for Microspheres with Ratiometric Barcode Lanthanide Encoding (MRBLEs)

**DOI:** 10.1101/402560

**Authors:** Björn Harink, Huy Nguyen, Kurt Thorn, Polly Fordyce

## Abstract

Multiplexed bioassays, in which multiple analytes of interest are probed in parallel within a single small volume, have greatly accelerated the pace of biological discovery. Bead-based multiplexed bioassays have many technical advantages, including near solution-phase kinetics, small sample volume requirements, many within-assay replicates to reduce measurement error, and, for some bead materials, the ability to synthesize analytes directly on beads via solid-phase synthesis. To allow bead-based multiplexing, analytes can be synthesized on spectrally encoded beads with a 1:1 linkage between analyte identity and embedded codes. Bead-bound analyte libraries can then be pooled and incubated with a fluorescently-labeled macromolecule of interest, allowing downstream quantification of interactions between the macromolecule and all analytes simultaneously via imaging alone. Extracting quantitative binding data from these images poses several computational image processing challenges, requiring the ability to identify all beads in each image, quantify bound fluorescent material associated with each bead, and determine their embedded spectral code to reveal analyte identities. Here, we present a novel open-source Python software package (the *mrbles* analysis package) that provides the necessary tools to: (1) find encoded beads in a bright-field microscopy image; (2) quantify bound fluorescent material associated with bead perimeters; (3) identify embedded ratiometric spectral codes within beads; and (4) return data aggregated by embedded code and for each individual bead. We demonstrate the utility of this package by applying it towards analyzing data generated via multiplexed measurement of calcineurin protein binding to MRBLEs (<u>M</u>icrospheres with <u>R</u>atiometric <u>B</u>arcode <u>L</u>anthanide <u>E</u>ncoding) containing known and mutant binding peptide motifs. We anticipate that this flexible package should be applicable to a wide variety of assays, including simple bead or droplet finding analysis, quantification of binding to non-encoded beads, and analysis of multiplexed assays that use ratiometric, spectrally encoded beads.

## Introduction

Multiplexing technologies enable the measurement of many analytes of interest within a single small volume, thereby saving precious reagents and reducing the cost and labor associated with each assay. Multiplexed bead-based assays, in particular, have been used for a wide range of applications in both research and diagnostics [1–8]. In these assays, different analytes are tethered to beads with a one-to-one linkage between analyte and beads. Bead-bound analyte libraries can then be pooled and tested in parallel for reaction with a probe molecule (*e.g.* testing protein binding or oligonucleotide hybridization to bead-bound peptides or oligos). Beads provide a support for solid-phase chemical synthesis (*e.g.* peptide or oligonucleotide synthesis), enabling rapid, precise, and low-cost library generation [9]. In addition, bead-based assays allow near-solution phase kinetics for measurement of up to hundreds of analytes simultaneously in volumes as low as 20 μL, with similar or better sensitivity and specificity relative to ELISA [1,10,11]. Finally, each assay can include many bead replicates for a particular analyte, reducing measurement error and enhancing the ability to detect subtle quantitative differences between analytes.

Multiplexing bead-based assays requires the ability to reliably identify all bead-bound analytes. This can be accomplished via a ‘one-code-one-compound’ approach in which beads are encoded and each analyte is tethered to or synthesized on beads bearing a different code (Fig. 1A) [9]. Various encoding strategies exist, including chemical encoding (in which beads are tagged with distinct chemical compounds) [1] and optical encoding (in which beads are produced with multiple shapes, embedded colors, or both) [12]. Optical encoding allows bead codes to be identified via imaging without a need for equipment-intensive downstream analysis (*e.g.* mass spectrometry or sequencing). In addition, optical encoding allows nondestructive code identification, facilitating kinetic monitoring of on-bead activity throughout an experiment. Finally, optically-encoded beads can be imaged with relatively cheap, or even mobile, equipment such as low-cost hyperspectral cameras, potentially enabling point-of-care applications [13].

**Fig 1.**
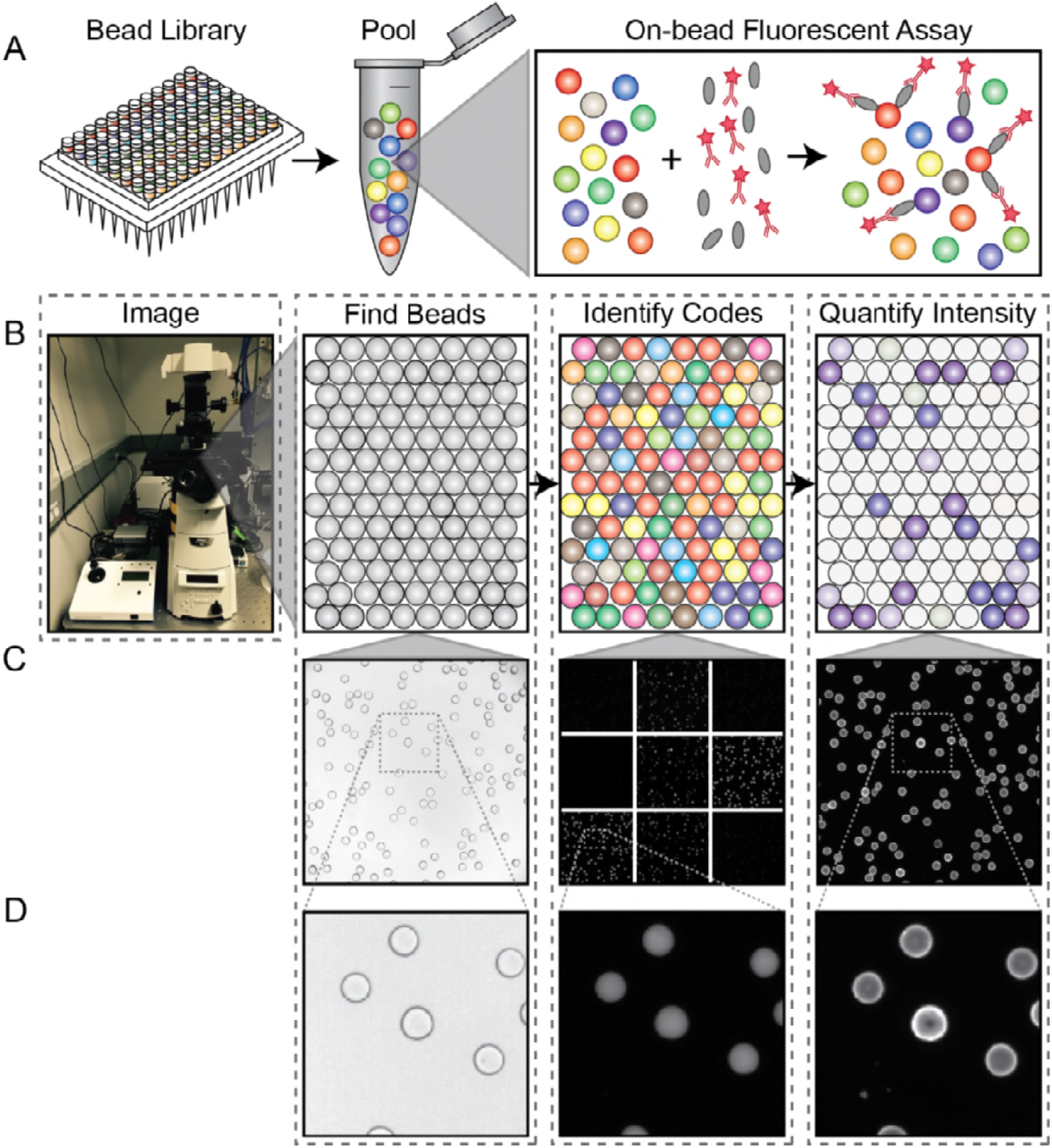
Overview of the experimental and analysis pipeline. **A)** Diagram of the MRBLE library assay. **B)** Diagram of the image and data analysis pipeline. **C)** From left to right: photograph depicting the quartz slide with the MBRLEs positioned on the microscope; bright-field image of MBRLEs; images of the 9 emission channels for decoding; image of the Cy5 channel used for assay quantification. **D)** Close-ups of images above, with the third image a close-up of one emission channel 6 (620 nm), which corresponds to a specific peak in Europium.

We recently developed a novel technology for producing spectrally encoded beads, which we term MRBLEs (<u>M</u>icrospheres with <u>R</u>atiometric <u>B</u>arcode <u>L</u>anthanide <u>E</u>ncoding). In this approach, microspheres are encoded via the ratiometric incorporation of lanthanide nanophosphors (LNP) particles. Compared to organic fluorophores or quantum dots, LNPs have very narrow emission spectra, making it possible to mix different LNPs together at multiple ratios and accurately deconvolve the spectral contributions from each species to ‘read’ codes. Multiple LNPs can be excited at a single wavelength that is spectrally orthogonal to wavelengths used to excite organic fluorophores, preserving the ability to detect up to 3 different types of fluorescently-labeled molecules bound to encoded beads in a single assay [14]. Using downconverting LNPs, we recently demonstrated the ability to produce and resolve MRBLEs containing > 1,100 distinct spectral codes [15]; even larger code spaces can be generated by including additional upconverting LNPs [16].

Accessing the large code spaces possible with LNP-based encoding requires the use of an image-based readout to collect sufficient photons from each bead to overcome shot noise, as LNP emission lifetimes are on the order of µs to ms (more than ˜ 1,000-fold slower than fluorescence emission) [17]. Identifying analytes and measuring binding therefore requires the development of image processing software capable of identifying individual beads, quantifying fluorescent material bound to each bead, and calculating ratios of embedded LNPs to identify the bead code and thus, the identity of the bound analyte (Fig. 1B). Identifying bead codes and quantifying binding additionally requires segmentation of each bead into an outer shell and an inner core, as LNPs that comprise the embedded spectral codes are located within the central core, while bound probes produce a ring of fluorescence around the outer bead margins (Fig. 1C,D). While multiple commercial software packages exist for decoding ratiometric spectral codes (*e.g.* BD FCAP, Illumina xPONENT, Bio-Rad Bio-Plex), these are optimized for flow cytometry data rather than images and are therefore incompatible with the use of LNPs. In addition, these commercial packages are closed-source, preventing critical modifications required for development of new assays.

Here, we present *mrbles*, an open-source software package for analysis of images acquired from multiplexed bioassays using spectrally encoded beads. This software is written in Python, a widely-used open-source programming language, and accompanied by an example Jupyter Notebook to be accessible even to users with limited programming experience. While we present this package as a method for decoding MRBLEs, *mrbles* should be broadly applicable to decoding and quantifying binding to unencoded beads as well as beads embedded with any ratiometric codes, including those based on organic fluorophores or quantum dots. Therefore, the analysis can be used for a wide range of potential applications, from imaging bead-bound libraries incubated with different concentrations of fluorescently-labeled macromolecules to extract binding affinities in high-throughput to imaging bead-bound libraries over time to extract binding kinetics. Unlike commercial software packages, which require images and inputs generated from proprietary commercial hardware, the *mrbles* package is compatible with a wide variety of inputs.

## Results

*The mrbles* analysis package is designed to automate image analysis and data extraction for multiplexed bioassays and to provide subsequent quality control, data analysis, and visualization tools. The software is set up in a modular fashion to provide data from each step in the pipeline, thereby enhancing generalizability (Fig. 2).

**Fig 2.**
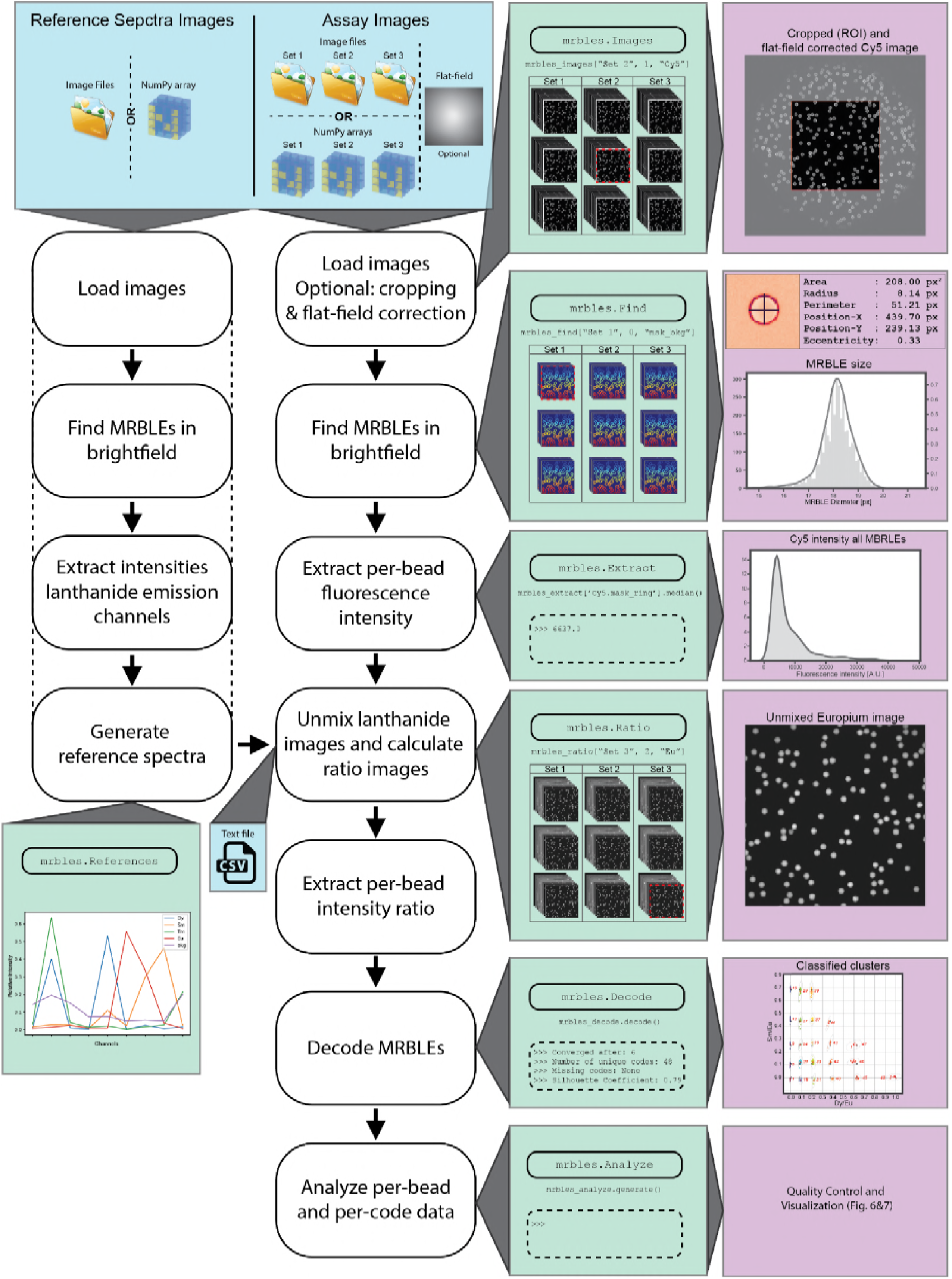
Diagram of analysis pipeline. Depicted are the top-level classes of the pipeline module, the required input data, and the various data output steps.

### Overview of the *mrbles* package architecture

The *mrbles* package contains four main modules (files): <u>data</u>, for the data structure classes and methods; <u>core</u>, for the core functionality and algorithms; <u>report</u>, for the quality control reports; and <u>pipeline</u>, for the front end of the analysis package. The analysis pipeline includes the following classes for step-by-step analysis (as diagrammed in Fig. 2):

1) *mrbles.Images*, for loading multiple image sets;

2) *mrbles.Find*, for finding beads and segmenting them into regions;

3) *mrbles.References*, for creating the reference spectra;

4) *mrbles.Ratio*, for generating spectrally unmixed and ratiometric images in individual coding channels;

5) *mrbles.Extract*, for per-MRBLE intensity extraction using regions from *mrbles.Find* and unmixed and ratiometric images from *mrbles.Ratio*;

6) *mrbles.Decode*, for identifying the code within each MRBLE; and

7) *mrbles.Analyze*, for per-code statistical parameters and generation of per-bead quality control reports.

### Loading multidimensional images via *mrbles.Images*

To begin any bead analysis, acquired images must be loaded into Python. Although all assays will likely contain images acquired at multiple slide locations (to capture information about all beads in the assay), the number of channels and number of images acquired at each location will vary based on the hardware used and the type of assay performed. For example, simple bead finding from bright-field images requires only a single image at each location, while full analysis of analyte binding to spectrally encoded beads requires a bright-field image (for bead finding), a variable number of images to identify embedded spectral codes (˜ 9 LNP emission images for decoding MRBLEs), and a variable number of fluorescence images (1-3, depending on the number of fluorophores used to detect bound material).

The *mrbles.Images* class provides a flexible method for handling the import of these various image formats for downstream analysis. For analyzing MRBLEs, this class can import OME-TIFF hyperstacks acquired using µManager [18] and convert them into a *mrbles* data frame for use throughout the rest of the pipeline. Using a user-provided folder and file pattern defined with regular expressions (regex), the class loads all images within the folder and extracts image metadata to allow specific images to be called later by file number and channel name using standard syntax (e.g. images[1, “Bright-field”]).

In the absence of OME-TIFF hyperstack images (*e.g*. if users only have a single image for bead finding from bright-field images, or users acquired images with a different acquisition software), the *mrbles.Images* class can alternately load user-provided multidimensional NumPy arrays in a Python dictionary with an optional list of names for each channel.

Loading images takes several minutes, depending on the amount of image data and the hardware used. To streamline image analysis and allow efficient processing of data sets containing multiple measurement conditions, multiple image sets can be loaded. Additional information about data input, required syntax, additional methods, and examples for *mrbles.Images* is available within the class documentation and the Github repository.

### Identifying and segmenting MRBLEs via *mrbles.Find*

Extracting quantitative information from individual beads requires the ability to reliably identify beads, even when packed together, while excluding other small contaminants (*e.g.* dust particles). Under bright-field illumination, MRBLEs appear as dark rings visible within a bright background (Figs. 3A,B), allowing bead identification via a simple thresholding procedure in which all pixels that exceed a particular value are used to define bead margins. Unlike traditional circle finding methods (*e.g.* Hough transform), this preserves morphological information about each MRBLE that can be used in downstream filtering to eliminate unwanted particles. In addition, the use of bright-field images to identify beads renders the *mrbles* package compatible with assays imaged with low-cost bright-field illumination.

**Fig 3.**
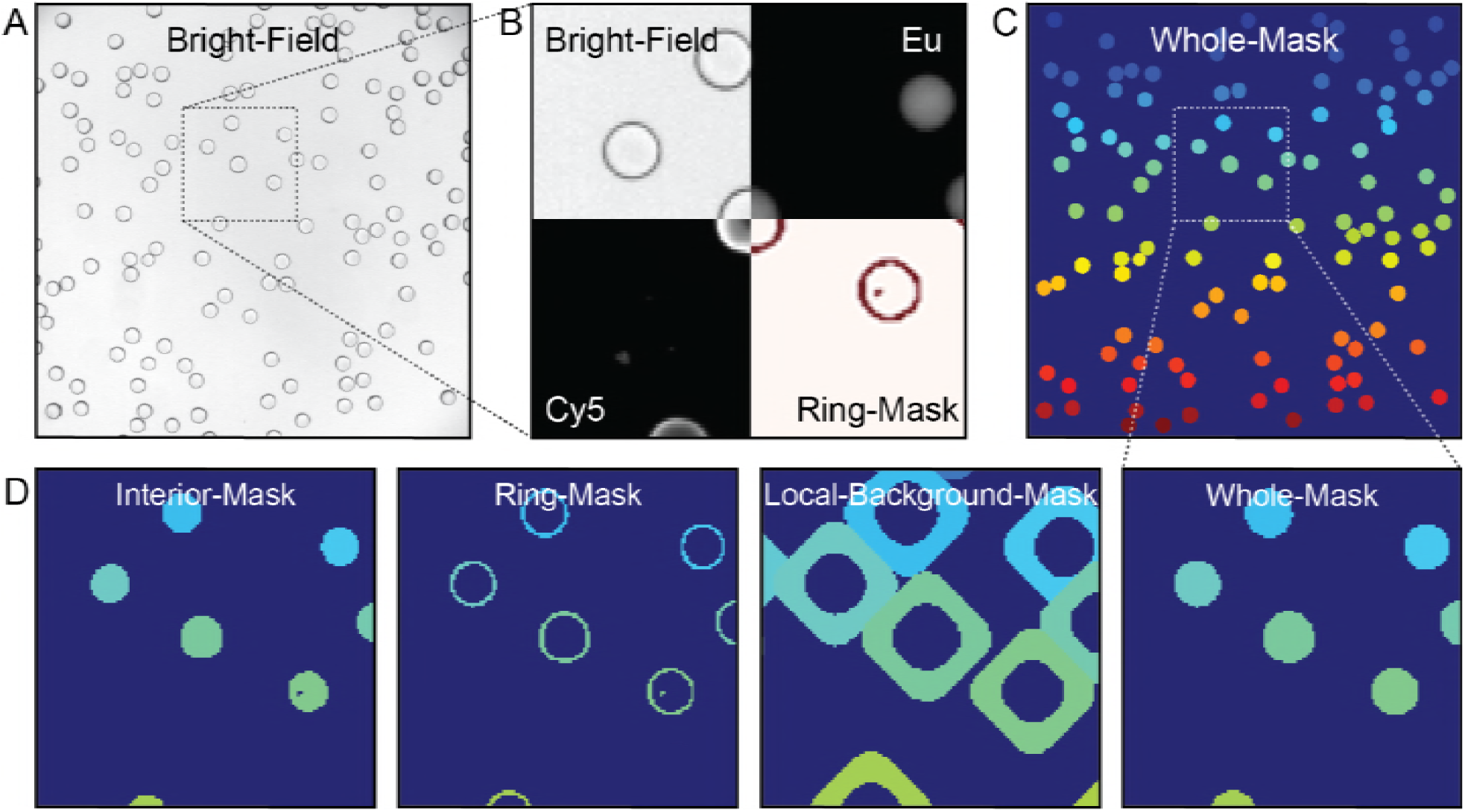
Images depicting the overlay of the assay region and various channels. **A)** bright-field image used for find the beads; **B)** cut-outs of fluorescent (Cy5) channel, bright-field, found ring region, and unmixed Europium channel; **C)** final mask depicting the whole region of each individual bead; **D)** zoomed-in region of the different mask regions generated from the mrbles.Find class. The color (a unique number) indicates the distinct region for one particular MRBLE. This color matches among the different masks and is unique to each identified MRBLE in these images. Dark blue represents the background.

Successfully delineating bead margins via intensity thresholding requires an appropriate threshold value, which can vary across the image due to local differences in illumination intensity. The *mrbles* package accounts for any local heterogeneity through the use of an adaptive Gaussian thresholding algorithm [19]. All pixels exceeding this threshold are used to create a binary mask that simultaneously identifies all beads in an image and identifies: (1) all pixels associated with that bead, (2) pixels defining an outer bead ring (representing the bead shell), (3) pixels fully enclosed by this ring (representing the bead interior), and (4) pixels just outside this ring (representing the local background) (Fig. 3B-D). To exclude dust, air bubbles, and other non-MRBLE particles from downstream analysis, *mrbles.Images* allows filtering of all found objects based on the total area and eccentricity.

Imaging densely packed bead samples improves overall assay throughput but poses additional image processing challenges. Beads in close proximity to one another have contiguous edges, yet pixels must be uniquely assigned to a single bead for downstream analysis. To do this, *mrbles.Find* employs a watershed algorithm that begins with the set of ‘interior’ pixels and then expands outward pixel-by-pixel until it touches either the edge of the identified bead ring or touches another assigned expanded bead region. At this point, the expansion stops, ensuring that closely packed beads are properly segmented from one another and each pixel is uniquely assigned to a single bead. After bead identification, all beads within each image are given a tracking number according to their location. This number, in combination with the image file number, allows unique tracking of each identified bead and all bead-associated data throughout the analysis process.

For experiments seeking to enumerate beads in bright-field images and quantify their size and geometry, the pipeline can be stopped at this point. Each bead object contains a variety of useful information, including: (1) pixels associated with each of 4 bead regions (whole, core, ring, and background) (2) bead centroid location, and (3) bead morphology information (*e.g.* eccentricity, area, diameter). The associated pixel regions can be inspected by viewing the mask ‘images’ (Fig. 3D) and are accessed using the same standard syntax as mentioned in the previous section (e.g. find_beads[‘Set A’, 3, ‘mask_ring’]). All numerical (tabular) centroid and morphology data is stored in a Pandas dataframe, which can be used for further analysis within the Jupyter notebook or output to standard text file format (*e.g.* csv, see Pandas package for documentation and *mrbles* GitHub repository for examples).

### Quantifying bound fluorescent material via *mrbles.Extract*

In bead-based binding assays, the strength of binding is determined by quantifying the intensity of fluorescently-labeled material bound to each bead [14,15]. The effective pore size within polymerized MRBLE hydrogels is small, sterically excluding large fluorescently-labeled probes (*e.g.* antibodies, proteins, and oligonucleotides). As a result, bound fluorescent proteins or oligonucleotides appear as a thin bright ring around the perimeter (Fig 3B). In MRBLE assays, these bright fluorescent pixels are the same as those delineating bead margins in the bright-field channel (Fig 3B), allowing bound fluorescence to be quantified by calculating the median fluorescence intensity for all pixels in the outer bead ‘ring’ segment identified by *mrbles.Find*.

The ultimate dynamic range and sensitivity of the assay depends on the signal-to-noise ratio of this median fluorescence intensity relative to the surrounding background. Fluorescence excitation using an LED or lamp-based light source can result in large spatial variations in the intensity of excitation across a sample field of view [20,21]. The *mrbles.Images* and *mrbles.Extract* classes allow normalization of these spatial variations by dividing experimental images by a flat-field ‘correction’ image of a highly concentrated dye sample [20,21] and using these corrected images to normalize fluorescence intensities across regions. The *mrbles.Images* class also allows users to specify a particular cropped region of interest (ROI) within each image. While a flat-field correction can compensate for systematic position-specific differences in excitation or collection efficiency, other experimental factors (*e.g.* dust, reflections, or slight tilting of the slide) can also skew local fluorescence signals. To account for these local variations and enhance discrimination of weak binding from background, *mrbles.Extract* performs a local background subtraction using the background region defined for each bead in *mrbles.Find.*

The fluorescence quantification portion of the analysis adds several additional pieces of information to each previously defined bead object, including median unprocessed, background, and background-subtracted fluorescence intensities for each given fluorescence channel.

### Calculating reference spectra for use in linear unmixing using *mrbles.References*

Isotropic spectral encoding can yield an extremely large code space, given by first approximation as *N = I*^*C*^, where *N* is the number of distinct spectral codes, *C* is the number of distinct coding species, and *I* is the number of intensity levels per coding species. Ratiometric encoding, in which one coding species is incorporated into all encoded particles, can enhance resolution of intensity levels by normalizing for small spatial differences in excitation intensity or photon collection efficiency. MRBLE spectral codes are generated by isotropically embedding multiple LNP species at different ratios within each bead, with one LNP species used as an internal fixed reference standard. To ‘read’ codes, measured intensities for each pixel across all channels are compared to ‘reference’ spectra for individual LNPs alone and a linear unmixing algorithm is used to identify the linear combination of LNPs most likely to have produced the observed pattern [22]. The *mrbles.References* class provides the code required to calculate or load these ‘reference’ spectra.

For MRBLE analysis, reference spectra are typically generated by synthesizing MRBLEs containing the maximum possible amount of each LNP alone and imaging them under standard experimental imaging conditions. The *mrbles.References* class loads these image stacks, finds beads within the bright-field images and segments each bead into regions (as described above), calculates the median pixel intensities within the ‘core’ of all found MRBLEs at each luminescent emission wavelength (435, 474, 536, 546, 572, 620, 630, 650, and 780 nm, under 292 nm deep UV excitation) [15], and stores these data in a Pandas dataframe. This analysis can be adapted to other spectrally encoded beads by using custom channels for each reference. In the event that reference bead images are not available, the *mrbles.References* class can also accept a delimited text file containing median intensities at each wavelength derived from fluorimetry or other methods.

MRBLE intensities observed under experimental conditions include contributions from LNPs as well as autofluorescence from slides, spectral bleed-through from fluorescence channels, and other residual background signals. Explicitly considering these contributions during linear unmixing improves the overall accuracy and confidence with which a bead can be assigned to a particular code. To generate this ‘background’ reference spectrum, the *mrbles.References* class calculates the median intensity of a user-defined region lacking beads from an experimental image across all LNP emission channels.

Finally, as a quality control, the *mrbles.References* can plot the reference spectra, the region used in the background spectrum, and store the reference spectra values in a text file (from the Pandas dataframe).

### Unmixing MRBLEs LNP emission intensities and calculate LNP ratio images using *mrbles.Ratio*

To calculate per-bead LNP ratios and identify embedded codes, measured intensity profiles must first be converted from luminescent channel intensities (Fig. 4A) into LNP channel contributions (Fig. 5B). To do this, the *mrbles.Ratio* class linearly unmixes median per-pixel LNP emission intensities relative to the LNP ‘reference’ spectra calculated above [23]. The *mrbles.Ratio* unmixes the per-pixel intensity profiles of each image in each LNP emission channel into probable contributions from each LNP by calculating the least-squares solution to a linear matrix equation:

**Fig 4.**
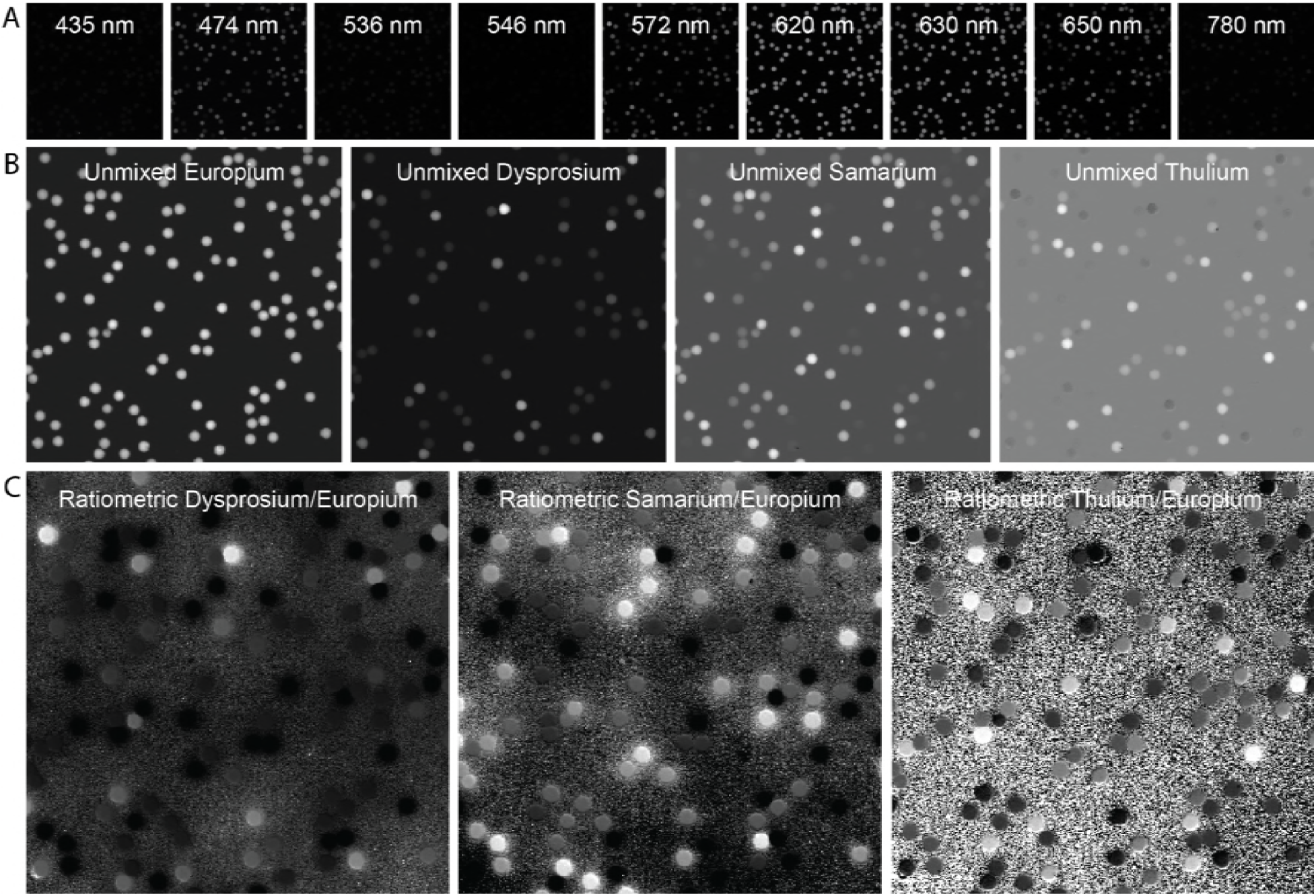
A) Original LNP emission channel images. **B)** Linear unmixed images using images from the LNP emission channels above. **C)** Ratiometric images resulting from division with the reference unmixed LNP (Europium).

**Fig 5.**
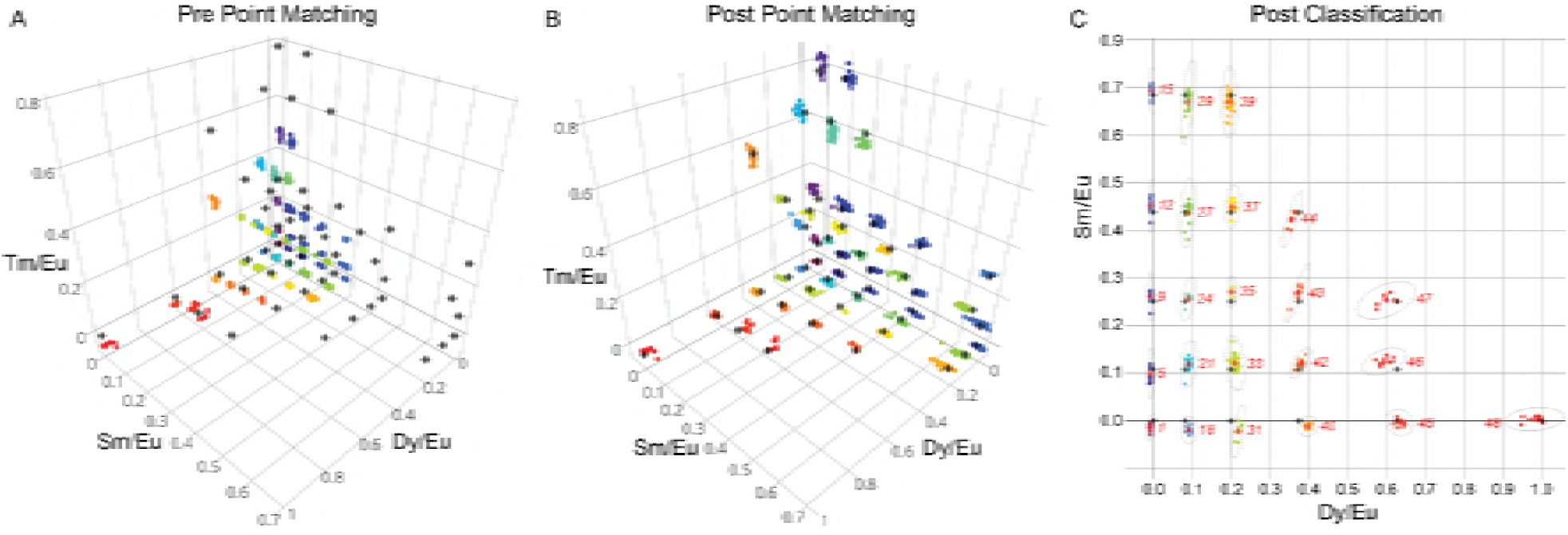
Code-ratio clusters prior and post alignment. Grey points are target points. Colored points are identified clusters. **A)** 3D scatter graph with clusters before point matching. **B)** 3D scatter graph with clusters after point matching, using Iterative Closest Point matching. **C)** 2D plot depicting post-classification code clustering and the per cluster 95%confidence interval (dotted line).

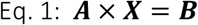

In this equation, ***A*** represents the ‘reference spectrum’ matrix for all LNP species; ***B*** contains pixel intensity information for all channels at a given location; and ***X*** represents the eigenvector solutions (linear weights reflecting likely contributions of each LNP species) for each pixel location. The *mrbles.Ratio* class then uses these weights to construct pseudo-intensity images for each LNP ‘channel’ (Fig. 4B). As a final step, the class normalizes the images for each variable LNP channel by the intensity of each pixel in the invariant ‘reference’ LNP channel, yielding ratiometric images (Fig. 4C). All of these images can be examined using the same syntax described in the image loading section (*e.g.* images[‘Set B’, 2, ‘Tm’]), making it possible to visually inspect the unmixed and ratio images.

### Identifying embedded codes within each MRBLE via *mrbles.Extract* and *mrbles.Decode*

To calculate LNP ratios associated with each bead, the *mrbles.Extract* class: (1) identifies all pixels in the linearly unmixed and ratiometric images associated with each bead core, and then (2) calculates median LNP levels and ratios for all of these pixels. To remove objects that resemble beads under bright-field imaging but are not true encoded particles (*e.g.* air bubbles), *mrbles.Extract* can filter out particles with invariant LNP levels and unmixed background levels that deviate significantly from the calculated mean value (using a user-defined threshold with a default value of > 2 standard deviations). This strategy ensures that all identified beads used in downstream analysis are actual MRBLEs with an accuracy of > 99.9% [15].

Determining the identity of the analyte associated with each bead requires determining the most probable code assignment for a given observed LNP ratio. However, while cluster assignment is trivial in many cases, it can be complicated by situations in which individual clusters are missing or measured levels appear scaled from expected reference levels. Spectral codes are typically generated according to a desired target matrix, providing information that can be leveraged to improve code assignment. For robust matching of clusters with their appropriate codes, the *mrbles.Decode* class stretches, scales, and rotates the overall cluster pattern to match the original designed code target ratios by implementing an iterative closest point (ICP) matching algorithm [24]. Commonly used in reconstructing 3D surfaces from different scans, such as Computed Tomography (CT), MRI, and optical scans for surgery [25,26], this algorithm minimizes the distances between two clouds of points or between the centroid positions of each cluster in the cloud and the nearest single points (in this case, the ratio cluster centroids and the target code ratios). In the final step, the *mrbles.Decode* class assigns ICP-transformed LNP ratio clusters (corresponding to unique MRBLE codes) to target code ratios using a Gaussian Mixture Model (GMM, scikit-learn) for supervised classification. Unlike *k*-means clustering, classification using a GMM is compatible with clusters with different variance and cluster sizes [27].

Visualizing the distribution of LNP ratios provides a convenient method to assess overall code quality, identify missing codes, and probe for spectral cross-talk between LNP species. After code assignment, the *mrbles.Decode* class can plot Dy/Ey, Sm/Eu, and Tm/Eu ratios for each bead in both 3D and 2D formats (Fig 5), and this function can be adapted for visualization of any 3 individual components of a coding scheme. Plotting the original LNP ratios can reveal, as mentioned before, scaling of the intended target ratios (Fig. 5A). In this case of Fig. 5A, there is a clear scaling of the Dy/Eu and Tm/Eu ratios caused by using reference beads containing different amounts of Dy and Tm than those used in the assay. These variations can also be caused by a change of experimental conditions (*e.g.* use of different buffer). Plotted transformed LNP ratios (post point matching) (Fig. 5B) for a typical MRBLEs code set reveal well-separated and easily visualized clusters that are unambiguously assigned to specific code clusters (Fig. 5C). As expected, the standard deviation of identified code clusters increases with the mean level of each included LNP, consistent with expectations for a largely Poisson encapsulation process. The quantitative relationship between cluster standard deviation and embedded LNP levels for each beads further provides critical information for choosing target code levels that discriminate clusters at a particular confidence level (*e.g.* 5 standard deviations).

The *mrbles.Decode* class outputs several parameters that indicate the quality of code assignment, including: (1) the number of steps required to converge the ICP; (2) the number of clusters (codes) found; (3) any missing clusters (codes), based on given target ratios; and (4) the silhouette score, an indicator of tightness of clustering for all clusters ranging from 0 (no clustering) to 1 (a singular point). Upon completion of this part of the pipeline, *mrbles.Decode* assigns each bead a code and provides a log probability and a probability to assess the classification for each individual bead. Beads with a low probability can be eliminated to remove ambiguously classified beads, potentially increasing possible code space significantly.

### Per-code data analysis summary via *mrbles.Analyze*

The ultimate goal of multiplexed binding measurements is to quantify interactions between an experimental probe of interest and a variety of different analytes. This, in turn, requires aggregating per-bead information to calculate binding statistics at the per-analyte (per-code) level. The *mrbles.Analyze* class returns a graphical report containing aggregated information for all beads, with the amount of information returned depending on which portions of the pipeline were run. All reports include a histogram of diameters for all identified beads from the *mrbles.Find* analysis, allowing calculation of median size and any observed size variability (Fig 6A).

**Fig 6.**
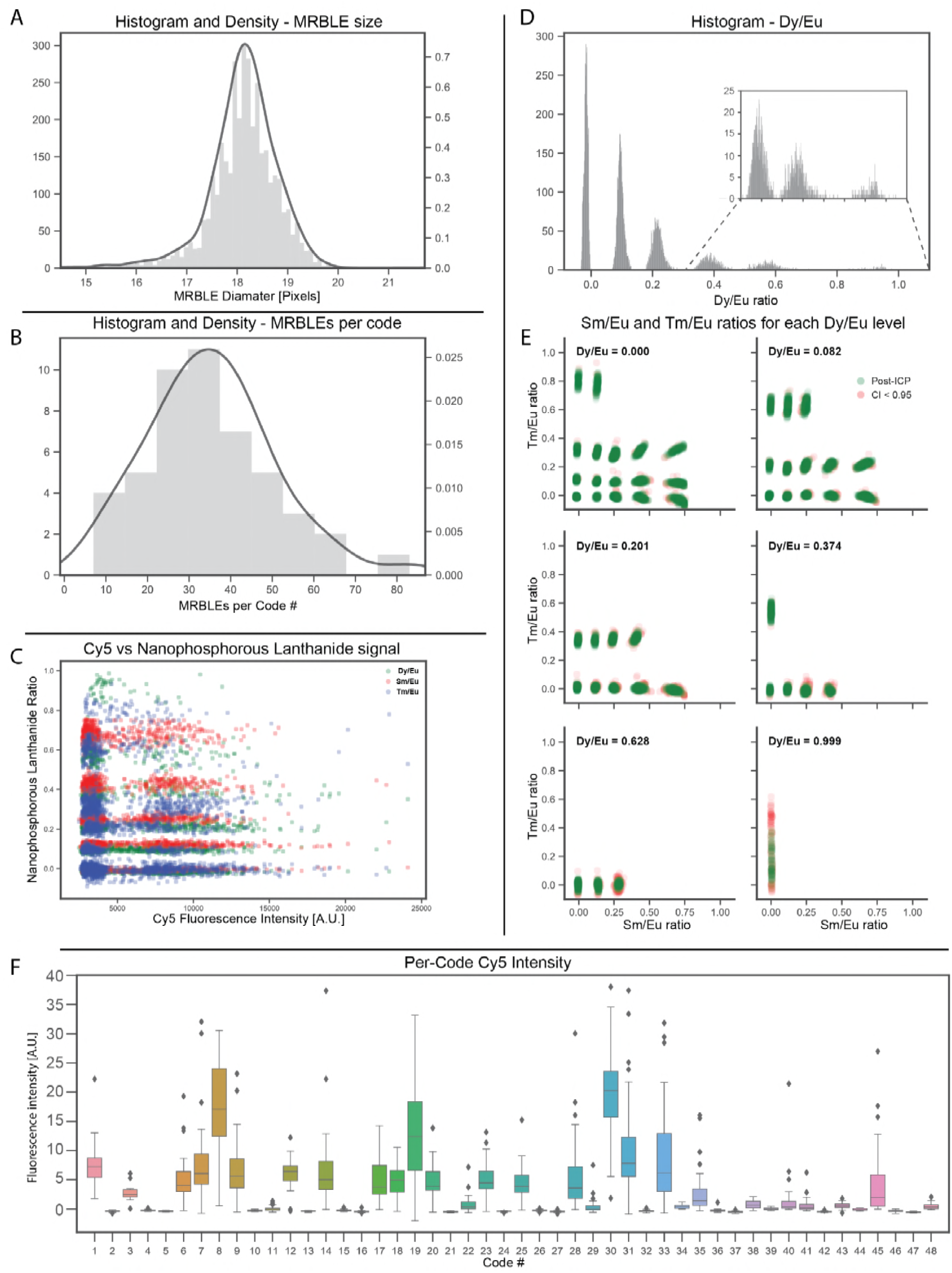
Quality control and final signal figures. **A)** MRBLE size distribution **B)** Per-code MRBLE distribution **C)** Fluorophore (Cy5) vs LNP signal, to demonstrate minimal correlation. **D)** Histogram of various Dy/Eu ratio levels **E)** The Sm/Eu and Tm/Eu ratio levels corresponding to the Dy/Eu ratios depicted above. **F)** Final per-code signal.

For analyses that proceeded through the *mrbles.Decode* class, the report produces a variety of additional graphs for experimental quality assessment and analysis. To reveal any heterogeneity in the number of beads synthesized or sampled, the report displays a histogram showing the number of beads identified per code (Fig. 6B). To probe for unwanted spectral cross-talk between LNP and fluorescence channels that could affect code calling or skew measured intensities, the report displays measured fluorescence intensity versus measured LNP ratios for each channel (Fig. 6C). To assess the quality and separability of observed code clusters for code sets with 3 ratiometric coding axes, the report also displays three-dimensional code clustering data in a two-dimensional format (Fig. 6D,E). First, measured intensities for a particular (user-specified) LNP ratio channel are binned and displayed as a histogram to reveal peaks associated with discrete ratiometric levels for that LNP (Fig. 6D). Next, the calculated ratios for the other two LNP channels for all beads associated one of these peaks are displayed as two-dimensional scatter (Fig. 6E). Finally, to quantify binding for each analyte in the assay, the report generates a box-and-whiskers plot displaying measured fluorescence intensities for each bead code (Fig 6F). Additional information about the identity of the analyte associated with each code can be included by loading a delimited text file, allowing generation of this box-and-whiskers plot as a function of analyte identity rather than bead code.

Along with this graphical report, the *mrbles.Analyze* class also returns a Pandas dataframe containing per-code statistical data for the given fluorescent channel (*e.g.* mean, median SD, etc.), providing users with information required to perform custom downstream analysis.

### Per-bead data analysis via *mrbles.Analyze*

Aggregated analysis of per-code data can reveal the presence of outliers, excessive variance, or other unexplained phenomena. Diagnosing the mechanisms responsible often requires the ability to visually inspect beads associated with a particular code. To facilitate this, the *mrbles.Analyze* class can also return a per-bead report that contains: (1) images of each bead under bright-field illumination, a user-specified fluorescence channel, and in unmixed LNP channels; (2) images of pixel regions associated with the whole bead, core region, ring region, and background region; and (3) intensity values and other quantitative parameters associated with that bead (*e.g.* fluorescence channel intensity, ratio intensity per LNP ratio image) (Fig 7). Bead reports can be generated for all beads in the assay (with returned results sorted by code number or image number), only beads associated with a particular code, or only beads associated with a particular image. Due to the large amount of data generated by this report (*e.g.* a typical assay has over 2,000 beads with 12 images per bead, creating 12,000 images for a full report), this function runs in several nested loops to optimize for speed and provides an estimated time before proceeding.

**Fig 7.**
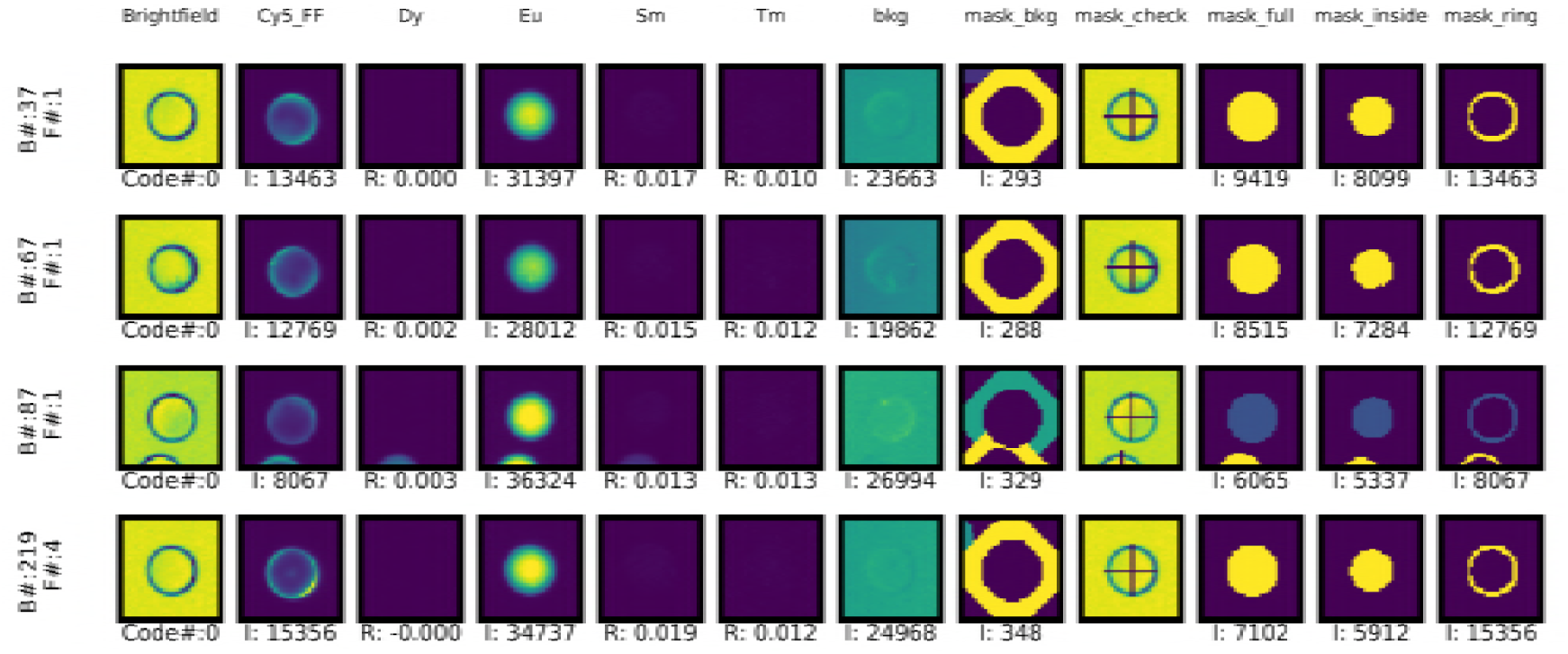
Example per-MRBLE image report. F=File number, I = Intensity in set region, and R = ratio.

## Discussion

The *mrbles* software package described here provides a flexible, open-source tool with broad applicability towards analyzing results from a variety of bead-based bioassays, including both standard and multiplexed assays on encoded beads. At the simplest level, the *mrbles.Find* class can be used to find beads in a set of bright-field images, facilitating fast and accurate determination of particle sizes and distributions for simple counting and aggregation experiments [28–30]. At the next level of complexity, the *mrbles.Extract* class in tandem with the *mrbles.Find* class allows accurate bead segmentation and quantification of fluorescence for macromolecules that appear as a thin ring around bead perimeters, including antibody-antigen interactions [5], DNA binding interactions [7], and protein-peptide interactions [14]. Finally, adding the *mrbles.References* and *mrbles.Decode* classes make it possible to reliably identify spectral codes associated with beads encoded via ratiometric encoding, including those produced via ‘on-the-fly’ optical encoding [31] or Raman-based encoding [32]. In all cases, the *mrbles.Analyze* class returns valuable information at both the per-code and per-bead levels via creation of graphical reports and output of simple delimited text files. Finally, the open-source nature of this software package enables collective efforts to optimize or modify it for multiplexed bead assays or other future purposes.

## Methods

### *mrbles* package installation and use

The *mrbles* package is available within the Python Packaging Index (PyPI) environment, allowing installation via the simple command “*pip install mrbles*“. It can then be imported as a module into any environment (including the command line, scripts, or a Jupyter notebook) using the command “*import mrbles*”. All source code is available through GitHub: https://github.com/FordyceLab/MRBLEs.

For demonstration and troubleshooting, the software package comes with a set of images from a MRBLEs assay measuring interactions between a fluorescently-labeled (Cy5) protein and peptides on different encoded beads. The package also includes example images for calculating individual LNP reference spectra, and a flat-field image for use in flat-field correcting measured Cy5 intensities. The GitHub repository also provides example Jupyter Notebook files analyzing these images with extensive explanations provided for each pipeline step described in this paper, following the structure of the software depicted in Fig 2. The source code is documented following Python NumPy docstring convention, giving users instant access to information to which parameters to provide and which methods are available. This documentation is also available on the GitHub Pages: https://fordycelab.github.io/MRBLEs.

### *mrbles* package module and folder structure

The *mrbles* package is structured in four main modules (files): <u>data</u>, for the data structure classes and methods; <u>core</u>, for the core functionality and algorithms; <u>report</u>, for the quality control reports; and <u>pipeline</u>, for the front end of the analysis package. The pipeline uses the classes and methods of the <u>core</u>, <u>report</u>, and <u>data</u> modules and can be called at the root of the package (e.g. *mrbles.Images*). All classes and methods in the other modules are preceded by their module name (e.g. *mrbles.data.ImageDataFrame*). These module files can be found in the “mrbles*”* folder in the source code available on GitHub. Additionally, the repository has an “examples” folder holding Jupyter Notebook examples; a “docs” folder for documentation generated from the inline docstrings; and a “data” folder containing example assay images, example reference images, a flat-field image, and input information files.

### MRBLEs imaging

For imaging, MRBLEs were loaded on a quartz glass slide (Electron Microscopy Sciences, 75 mm × 25 mm, 1 mm thick, cat. # 72250-03), covered with a quartz coverslip (Electron Microscopy Sciences, 25 mm × 25 mm, 0.2 mm thick, cat. # 72250-02), and placed on a Nikon inverted Ti-E microscope for imaging. Deep UV and bright-field illumination were provided by a Xenon arc lamp equipped with a 292/27 nm bandpass filter (Semrock) and a 409 nm longpass filter (Semrock), for UV and bright-field respectively, directed through a UV-extended range liquid light guide (Sutter) mounted via a custom 3D printed assembly in place of the microscope condenser above the sample. An additional longpass UV-rejection OD2 filter (Edmund Optics, cat. # 49024) was placed on top of the objective (below the sample). Fluorescence excitation was provided by a SOLA light engine (Lumencor) directed through a variety of standard fluorescence filter cubes (Semrock) mounted in a motorized filter turret. To image LNP emission, light emitted by the sample was passed through multiple filters (435/40, 474/10, 536/40, 546/6, 572/15, 620/14, 630/92, 650/13, and 780/25, Semrock) mounted in a motorized filter wheel (Sutter Instruments) and imaged on an sCMOS camera (Andor). All images were acquired at 2x binning using a 4x objective (Nikon CFI Plan Fluor) with exposure times per channel of 200, 450, 200, 200, 300, 200, 150, 350, 700 ms, respectively.

### MRBLEs assay

Images and data used in this paper are of a MRBLEs 48-plex peptide library composed of systematic variations within known calcineurin peptide substrates, and a scrambled peptide control. Peptides were attached to the MRBLE beads by solid state peptide synthesis, as described previously [15]. The MRBLE beads were passivated with blocking buffer (0.1% v/v TWEEN 20, 5% w/v bovine serum albumin, in 1X phosphate buffered saline, pH = 7.5, Sigma-Aldrich) overnight at 4 °C on a rotator. Afterwards, the blocking buffer was exchanged with binding buffer (50 mM Tris pH = 7.5, 150 mM NaCl, 0.1% v/v TWEEN 20, Sigma-Aldrich) with three cycles of pelleting, decanting, and resuspension. The experiment was performed with varying concentration of calcineurin containing a 6xHis-tag (expressed and purified [35]). Prior to incubation, calcineurin (2.5 µM) was labeled with anti-6xHis-DyLight-650-antibody (Abcam, ab117504) in a 1:1 ratio in binding buffer for 1 hour at 4 °C. For each concentration of CN:α-6xHis-Dy650 complex (100, 250, 500, and 1,000 nM) approximately 3,000 beads were incubated in a final volume of 100 µL. All the tubes were left on a rotator at 4 °C for ˜ 6 hours and then imaged after decanting and washing once with wash buffer (0.1% v/v TWEEN 20, 1X phosphate buffer saline, pH = 7.5).

### Flat field correction images

The flat-field image for channel was generated from taking images of a highly concentrated dye solution (for Cy5 or similar dyes: 100 mg·mL^−1^ Erioglaucine disodium salt in water, Sigma-Aldrich) with the same quartz slide and coverslip as used in the MRBLEs assay. The solution was sonicated for one hour and then filtered (0.20 µm, MilliPore) prior to use. The slide with 20 µL of dye solution was image in a tiled (9 × 9) arrangement with an overlap of 50%. Using Fiji [33], a median image was created from these 81 images. Additionally, a mean dark-field image was created from taking 500 images with a blanked camera to record the intrinsic noise pattern of the camera. This dark-field image was subtracted from the median dye solution image to produce a flat-field image [20,21].

### mrbles.Find

Imaging default settings were optimized for polyethylene glycol beads that subtend ˜ 10-20 pixels (35-70 µm beads). Current adaptive thresholding algorithm (opencv) default settings are set to a moving block size of 15 pixels, with a threshold of 11; the watershed algorithm uses the software package’s own default setting (scikit-image). Default settings for total area and eccentricity are set to 50-150% of accepted bead size and a maximum of 0.65, respectively. The bead finding processes is optimized for parallel computing (multiprocessing) to speed up the process. This option can be disabled, since this option is slower on computers with limited resources.

### mrbles.Ratio

In Eq. 1, ***A*** is a *c* × *r* matrix of intensities in which each row contains intensity information for a particular (LNP) emission channel (*c*), and each column contains the reference (*r*) intensities for a different (LNP) species and the background (*e.g.* 9 channels with 5 reference spectra: 9 × 5); ***B*** is a *c* × *p* matrix of observed intensities with each row containing the LNP channel intensity, and each column the pixel (*p*) intensity for each pixel in the corresponding (flattened) image (*e.g.* 9 channel images with each 96 × 96 = 9,216 pixels: 9 × 9,216); and ***X*** is a *r* × *p* matrix in which each row contains the contributions for each spectra most likely to have produced the observed pattern, and the columns the contribution for each pixel (*e.g.* combining mentioned example matrix shape: 5 × 9,216). These images are then reconstructed to its original image size from the input (*e.g.* 5 × 96 × 96), resulting in a pseudo-intensity image for each LNP ‘channel’ (Fig. 4B).

### mrbles.Decode

The ICP part of this class is a distance minimization function that uses a pairwise distance function (*scikit-learn*) to return the distance from all ratios from each bead to all target ratios. First, the target ratio closest to each bead’s ratio is chosen as its matched code ratio. Next, the least squares solution (*NumPy*) is used to find a new transformation matrix, where ***A*** is the distance matrix for all beads, ***B*** the matched code ratios for all beads, and ***X*** the eigenvector solutions for the transformation matrix. This transformation matrix can be divided into a scaling/rotation matrix (***T***) and the offset vector (***o***), which are used to calculate the cost function for distance minimization:

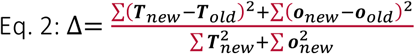

ICP is sensitive to outliers and overfitting, and therefore requires a proper initial scaling transformation and initial offset in order to find a global minimum. Empirical testing has revealed that generating an initial scaling transformation created by dividing the standard deviation (SD) of the ratios for each LNP dimension by the spread of the target ratio levels (SD) in each dimension works well as an initial transformation. This avoids an initial transformation based on outliers and correctly accounts for missing clusters on the outside of the ratio code space. To compensate for missing or shifted clusters or outliers within the ratio code space, the top percentile (typically 0.1-1%, adjustable with a default of 1%) of points with the furthest Euclidean distance from the target point is removed from the least squares fit at each iteration.

The final code classification is done by GMM (*scikit-learn*) in full variance mode, allowing variable variance for each LNP dimension, with supervised learning by using the target ratios (codes). Each code is given equal initial weighting, assuming there are approximately equal amounts of each code, and an initial covariance matrix (default sigma: 10^−5^). The GMM returns the assigned code for each bead and the log probability and probability of this assignment (using the LogSumExp algorithm, see *scikit-learn* documentation and [34]). Confidence interval ellipses for each cluster are calculated from the computed eigenvalues (magnitude in each direction, *e.g.* 1.96 × **v** × 2 for 95%) and eigenvectors (angle of ellipse) of the GMM covariance matrix (*numpy*) [35].

Author contributions
B.H, H.N., K.T., and P.M.F conceived of the software package. B.H. and H.N. developed the software package. K.T. and P.M.F. provided guidance during development. B.H. and P.M.F. wrote the manuscript.

## References

1. Elshal M, Mccoy J. Multiplex bead array assays: Performance evaluation and comparison of sensitivity to ELISA. Methods. 2006;38: 317–323.

2. Dincer C, Bruch R, Kling A, Dittrich PS, Urban GA. Multiplexed Point-of-Care Testing – xPOCT. Trends in Biotechnology. 2017;35: 728–742. doi:10.1016/j.tibtech.2017.03.01.

3. Tighe PJ, Ryder RR, Todd I, Fairclough LC. ELISA in the multiplex era: potentials and pitfalls. Proteomics Clin Appl. 2015;9: 406–422. doi:10.1002/prca.20140013.

4. Xiao Q, Luechapanichkul R, Zhai Y, Pei D. Specificity Profiling of Protein Phosphatases toward Phosphoseryl and Phosphothreonyl Peptides. J Am Chem Soc. 2013;135: 9760– 9767. doi:10.1021/ja401692.

5. Singhal A, Haynes CA, Hansen CL. Microfluidic Measurement of Antibody−Antigen Binding Kinetics from Low-Abundance Samples and Single Cells. Anal Chem. 2010;82: 8671–8679. doi:10.1021/ac101956.

6. Kang H, Jeong S, Koh Y, Cha MG, Yang J-K, Kyeong S, et al. Direct Identification of On-Bead Peptides Using Surface-Enhanced Raman Spectroscopic Barcoding System for High-Throughput Bioanalysis. Scientific Reports. 2015;5: 10144. doi:10.1038/srep1014.

7. Klamp T, Camps M, Nieto B, Guasch F, Ranasinghe RT, Wiedemann J, et al. Highly Rapid Amplification-Free and Quantitative DNA Imaging Assay. Scientific Reports. 2013;3: 1852. doi:10.1038/srep0185.

8. Subramanian A, Narayan R, Corsello SM, Peck DD, Natoli TE, Lu X, et al. A Next Generation Connectivity Map: L1000 Platform and the First 1,000,000 Profiles. Cell. 2017;171: 1437-1452.e17. doi:10.1016/j.cell.2017.10.04.

9. Liu R, Marik J, Lam KS. A Novel Peptide-Based Encoding System for “One-Bead One-Compound” Peptidomimetic and Small Molecule Combinatorial Libraries. J Am Chem Soc. 2002;124: 7678–7680.

10. Nolan JP, Mandy F. Multiplexed and microparticle-based analyses: Quantitative tools for the large-scale analysis of biological systems. Cytometry A. 2006;69A: 318–325.

11. Leng Y, Sun K, Chen X, Li W. Suspension arrays based on nanoparticle-encoded microspheres for high-throughput multiplexed detection. Chem Soc Rev. 2015;44: 5552– 5595.

12. Braeckmans K, De Smedt SC, Leblans M, Pauwels R, Demeester J. Encoding microcarriers: present and future technologies. Nat Rev Drug Discov. 2002;1: 447–456.

13. Habel R, Kudenov M, Wimmer M. Practical Spectral Photography. Comput Graph Forum. 2012;31: 449–458.

14. Nguyen HQ, Brower K, Harink B, Baxter B, Thorn KS, Fordyce PM. Peptide library synthesis on spectrally encoded beads for multiplexed protein/peptide bioassays. Microfluidics, BioMEMS, and Medical Microsystems XV. 2017.

15. Nguyen HQ, Baxter BC, Brower K, Diaz-Botia CA, DeRisi JL, Fordyce PM, et al. Programmable Microfluidic Synthesis of Over One Thousand Uniquely Identifiable Spectral Codes. Advanced Optical Materials. 2016;5: 1600548.

16. Boyer J-C, Vetrone F, Cuccia LA, Capobianco JA. Synthesis of Colloidal Upconverting NaYF4 Nanocrystals Doped with Er3+, Yb3+ and Tm3+, Yb3+ via Thermal Decomposition of Lanthanide Trifluoroacetate Precursors. J Am Chem Soc. 2006;128: 7444–7445. doi:10.1021/ja061848.

17. Singh NS, Ningthoujam RS, Luwang MN, Singh SD, Vatsa RK. Luminescence, lifetime and quantum yield studies of YVO4:Ln3+ (Ln3+=Dy3+, Eu3+) nanoparticles: Concentration and annealing effects. Chemical Physics Letters. 2009;480: 237–242. doi:10.1016/j.cplett.2009.09.00.

18. Edelstein AD, Tsuchida MA, Amodaj N, Pinkard H, Vale RD, Stuurman N. Advanced methods of microscope control using μManager software. Journal of Biological Methods. 2014;1: 10.

19. Gonzales RF, Woods RE. Digital Image Processing. 4th ed. London: Pearson; 2018.

20. Model MA. Intensity Calibration and Shading Correction for Fluorescence Microscopes. Current Protocols in Cytometry. 37: 10.14.1-10.14.7. doi:10.1002/0471142956.cy1014s3.

21. Model MA, Burkhardt JK. A standard for calibration and shading correction of a fluorescence microscope. Cytometry. 2001;44: 309–316.

22. Gerver RE, Gómez-Sjöberg R, Baxter BC, Thorn KS, Fordyce PM, Diaz-Botia CA, et al. Programmable microfluidic synthesis of spectrally encoded microspheres. Lab Chip. 2012;12: 4716–4723.

23. Keshava N, Mustard JF. Spectral unmixing. IEEE Signal Processing Magazine. 2002;19: 44–57. doi:10.1109/79.97472.

24. Besl PJ, McKay ND. A method for registration of 3-D shapes. IEEE Transactions on Pattern Analysis and Machine Intelligence. 1992;14: 239–256. doi:10.1109/34.12179.

25. Cutter JR, Styles IB, Leonardis A, Dehghani H. Image-based Registration for a Neurosurgical Robot: Comparison Using Iterative Closest Point and Coherent Point Drift Algorithms. Procedia Computer Science. 2016;90: 28–34. doi:10.1016/j.procs.2016.07.00.

26. Beek M, Small CF, Ellis RE, Sellens RW, Pichora DR. Bone alignment using the iterative closest point algorithm. J Appl Biomech. 2010;26: 526–530.

27. Su T, Dy JG. In search of deterministic methods for initializing K-means and Gaussian mixture clustering. Intelligent Data Analysis. 2007;11: 319–338.

28. Kemala T, Budianto E, Soegiyono B. Preparation and characterization of microspheres based on blend of poly(lactic acid) and poly(?-caprolactone) with poly(vinyl alcohol) as emulsifier. Arabian Journal of Chemistry. 2012;5: 103–108. doi:10.1016/j.arabjc.2010.08.00.

29. Rajan NK, Rajauria S, Ray T, Pennathur S, Cleland AN. A simple microfluidic aggregation analyzer for the specific, sensitive and multiplexed quantification of proteins in a serum environment. Biosensors and Bioelectronics. 2016;77: 1062–1069. doi:10.1016/j.bios.2015.10.09.

30. Wiklund M, Hertz HM. Ultrasonic enhancement of bead-based bioaffinity assays. Lab Chip. 2006;6: 1279–1292. doi:10.1039/B609184.

31. Marcon L, Battersby BJ, Rühmann A, Ford K, Daley M, Lawrie GA, et al. “On-the-fly” optical encoding of combinatorial peptide libraries for profiling of protease specificity. Mol Biosyst. 2010;6: 225–233.

32. Hu F, Zeng C, Long R, Miao Y, Wei L, Xu Q, et al. Supermultiplexed optical imaging and barcoding with engineered polyynes. Nat Methods. 2018;

33. Schindelin J, Arganda-Carreras I, Frise E, Kaynig V, Longair M, Pietzsch T, et al. Fiji: an open-source platform for biological-image analysis. Nature Methods. 2012;9: 676–682. doi:10.1038/nmeth.201.

34. Nielsen F, Sun K. Guaranteed bounds on the Kullback-Leibler divergence of univariate mixtures using piecewise log-sum-exp inequalities. Entropy. 2016;18: 442. doi:10.3390/e1812044.

35. Strang G. Determinants. Linear Algebra and Its Applications. 2nd ed. Orlando: American Press, Inc.; 1980. p. 222.

